# Optically pumped magnetometers disclose magnetic field components of the muscular action potential

**DOI:** 10.1101/2020.07.30.228882

**Authors:** Philip J. Broser, Thomas Middelmann, Davide Sometti, Christoph Braun

**Affiliations:** Children’s hospital of eastern Switzerland, Sankt Gallen, Switzerland; Physikalisch technische Bundesanstalt, Berlin, Germany; MEG Center, University of Tübingen, Germany; Hertie-Institute for Clinical Brain Research, Tübingen, Germany; CIMeC, Center for Mind/Brain Sciences, Tübingen, Germany

**Keywords:** peripheral nerve system, muscle action potential, optically pumped magnetometer, finite wire model, magnetic moving dipole model, magnetomyography (MMG), magnetoneurography (MNG)

## Abstract

**Aim:** To track the magnetic field generated by the propagating muscle action potential (MAP).

**Method:** In this prospective, proof of principle study, the magnetic activity of the intrinsic foot muscle after electric stimulation of the tibial nerve was measured using optically pumped magnetometers (OPMs). A classical biophysical electric dipole model of the propagating MAP was implemented to model the source of the data.

**Results:** The signal profile generated by the activity of the intrinsic foot muscles was measured by four OPM devices. Three devices were located above the same muscle to compare the direction and the strength of the magnetic signal while propagating along the muscle.

**Interpretation:** OPM devices allow for a new, non-invasive way to study MAP patterns. Since magnetic fields are less altered by the tissue surrounding the dipole source compared to electric activity, a precise analysis of the spatial characteristics and temporal dynamics of the MAP is possible. The classic electric dipole model explains major but not all aspects of the magnetic field. The field has longitudinal components generated by intrinsic structures of the muscle fibre. By understanding these magnetic components, new methods could be developed to analyse the muscular signal transduction pathway in greater detail.

**What this paper adds:**

- Technological concepts to record and analyse the small magnetic fields generated by electric muscular activity
- Model to link the signals measured by the OPM sensors to the underlying physiology
- Insights into the propagation of muscle action potential and the sequential control of motor activity

## 1 Introduction

The molecular control of the contractile apparatus of the skeletal muscle is critically dependent on the muscle action potential (MAP) propagating along the muscle fibre after initiation at the neuromuscular endplate (Moritani T, Stegeman D, Merletti R, 2004). After activation of the postsynaptic membrane by acetylcholine (AcH), several different positively charged ions can cross these channels (Martonosi, 2000). The influx of these ions depolarises the membrane of the muscle fibre, the voltage gated sodium channels open and a MAP is generated which propagates along the muscle fibre in both the proximal and distal directions (Farina and Merletti, 2004). In contrast to the neural action potential, the MAP is established by sodium, potassium and calcium ions (Martonosi, 2000). The speed of the MAP is an estimated 4–6 m/s (Farina and Merletti, 2004) slower than the action potentials of motor neurons (around 50 m/s, Ghezzi et al., 1991). Similar to the neuronal action potential (Wikswo et al., 1980), the MAP leads to a specific magnetic field (Zuo et al., 2020), which can be measured by superconducting quantum interference devices (SQUIDs, Reincke, 1993) or optically pumped magnetometers (OPMs) (Broser et al., 2018). However, there are significant anatomical differences between the axon of a neuron and a muscle fibre. First of all, the cross-sectional diameter of a muscle fibre is in the range of 20–100 μm, whereas an axon is typically 1 μm in diameter. Further, the membrane of a muscle fibre has a high number of cylindrical infoldings, known as T-tubuli. The MAP is propagated deep in the myocyte along the T-tubuli, in close proximity to the contractile apparatus. The MAP along the T-tubuli generates a radial current that creates an additional magnetic field component in the longitudinal direction of the fibre.

The magnetic fields generated by the MAP are in the range of 0.01 to 0.1 nanotesla (nT), and therefore, special devices are required to record these small magnetic fields. OPMs are newly developed sensors (Mhaskar et al., 2012; Colombo et al., 2016; Labyt et al., 2019) that can measure these small magnetic fields. Unlike SQUIDs (Reincke, 1993), OPMs are much smaller and can be placed in close proximity to the muscle.

In this study, we developed a measurement setup to record the MAP of the intrinsic foot muscles generated after electric stimulation of the tibial nerve using OPM devices. The magnetic field was measured in two orthogonal directions. The signal-to-noise ratio and reproducibility was tested. Given that magnetic field distributions are geometrically complex, and an intuitive guess of the source of the magnetic field might be misleading, we implemented a classical electric dipole model to explain the sources of the magnetic fields.

The model is based on an electric dipole propagating along the muscle fibre. The muscle fibre was approximated as a finite wire, and the MAP was modelled according to the study by Rosenfalck (1969).

Given that this model explains some but not all aspects of the magnetic field, a second model based on a moving magnetic dipole was in addition tested. This model is motivated by the estimation of the radial currents inside the muscle as found by Henneberg and Roberge (1997). In comparison to the first model, this model takes into consideration the radial currents resulting from the MAP propagating along the T-tubuli and thus generating a magnetic field component into the longitudinal direction of the muscle fibre.

## 2 METHOD

### 2.1 Recording of the magnetic activity of the MAP

A single-subject, proof of principal study was conducted at the MEG Center of the University of Tübingen in January 2020. The experiment was designed specifically to record the magnetic activity of the intrinsic foot muscles after tibial nerve stimulation.

The electric stimulus used to stimulate the nerve interferes with the OPM sensors for about 10 ms. Thus, the stimulation site was chosen such that the time delay between stimulation and muscle activity was greater than 10 ms. Therefore, the tibial nerve was stimulated at the level of the knee.

The study aimed to record the activity of the abductor hallucis brevis. This muscle is innervated by the tibial nerve (Figure 1A). The abductor hallucis brevis muscle is located at the medial border of the foot (Figure 1C) and was localised by palpation by an experienced clinical doctor (PB). The muscle inserts on the OS calcaneus. The muscle body is thick and flattens as it stretches forward to the big toe. The muscle ends in a common tendon with the flexor hallucis brevis. This tendon ends on the medial surface of the base of the first proximal phalanx (big toe, hallux). Additional intrinsic foot muscles are located close to the big toe (musculus flexor hallucis and musculus adductor hallucis). These muscles are also innervated by the tibial nerve, and therefore, their MAP could add signal components to the recording (Figure 1C).

**Figure 1:**
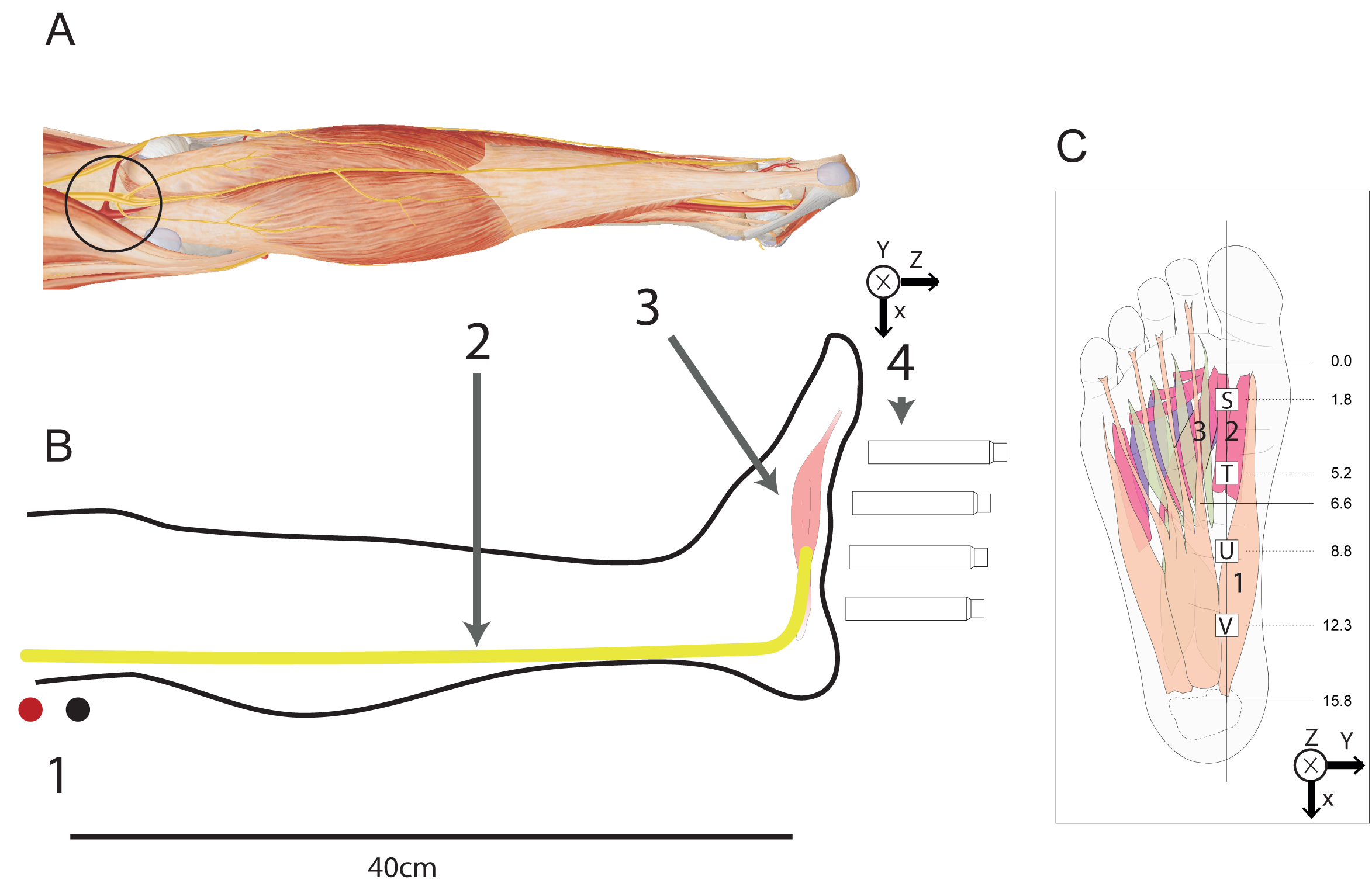
Principal setup of the experiment. (A) Schematic anatomical drawing of the lower leg. The black circle at the level of the knee highlights the area on the skin where the tibial nerve was stimulated. (B) Schematic drawing of the lower leg. Lateral view onto the medial aspect of the lower leg together with the relevant structures and technical equipment. (B1) Location of the self-adherent stimulation electrodes (black = cathode, red = anode). (B2) Tibial nerve. (B3) Intrinsic foot muscles, specifically the musculus abductor hallucis. (B4) Four OPM recording devices. Coordinate system in the upper right. The y axis points out of the drawing area. (C) Position of the four OPM devices (S, T, U and V) in relation to the anatomical structure. (C1) Musculus abductor hallucis. (C2) Musculus flexor hallucis brevis. (C3) Musculus adductor hallucis. The x axis points from the big toe to the heel. The y axis points from digit V to digit I.

During the experiment, the tibial nerve was stimulated by two electrodes placed on the skin on the dorsal side of the knee (Figure 1B) at a distance of about 40 cm proximal to the ankle. A series of four OPMs (labelled S, T, U and V) were positioned longitudinal to the medial border of the foot (Figure 1B). The OPM devices were placed such that the z-axis of the sensor pointed towards the plantar side of the foot, and the y axis pointed in the lateral direction (Figure 1B). The distance between neighbouring sensors was 3.5 cm. The sensors T, U and V were located above the musculus abductor hallucis and were therefore the main sensors of interest in the study. The participant (a co-author of this paper) gave his informed consent to participate in the study and agreed to the publication of his data. The study was performed in accordance with the Declaration of Helsinki (World Medical Association, 2001).

### 2.2 Experimental setup

The detailed experimental procedure (Figure 1A,B) began with the subject (a 30-year-old male) lying on a comfortable bed inside of a magnetic shielded room (Ak3b, VAC Vacuumschmelze, Darmstadt, Germany). The subject rested his right leg on a pillow. The right foot was placed in a cardboard box (25 × 15 cm) filled with rigid foam with a cavity to accommodate the foot. The bottom of the box had four slots to host the OPMs (QZFM-gen-1.5, QuSpin Inc., Louisville, CO, USA). These magnetometers were based on an optically detected zero-field resonance in hot rubidium vapor, which was contained in a vapor cell measuring 3 × 3 × 3 mm^3^. The centre of the cell had a distance of 6.2 mm to the exterior of the housing, which measured 13 × 19 × 85 mm^3^. This small size allows for easy handling of these sensors and flexibility to adapt the sensors to the specific geometrical situation (Alem et al., 2015; Boto et al., 2017, Osborne et al., 2018). The OPMs are capable of measuring two components of the magnetic field vector: the y and z direction (Figure 1B). They provided a magnetic field sensitivity in the order of 15 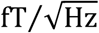 in a bandwidth of 3–135 Hz and a dynamic range of a few nT. To adapt to a non-zero magnetic background field, the sensors were equipped with internal compensation coils, which can cancel magnetic background fields with a strength of up to 200 nT in the sensing vapor cell.

Our OPM measurement system consisted of a total of five OPMs, which simultaneously recorded the magnetic field in two orthogonal directions (y and z). As described above, four sensors were placed inside the slots of the box along a line running across the medial plantar side of the foot (Figure 1C). A fifth sensor was placed just outside the box to measure the magnetic background activity.

In order to synchronise the muscular activity with the recording system, the MAP was recorded after electric stimulation of the nerve. To this end, self-adherent electrodes were placed on the skin of the subject in close proximity to the tibial nerve (Figure 1A,B) at the level of the knee. The stimulator (System Plus Fire, Micromed, Venice, Italy) was placed outside the magnetically shielded room, and the two-electrode cables (cathode and anode) were routed through small holes into the magnetic chamber. The tibial nerve was stimulated with a monopolar square wave pulse of 500 μs duration and an intensity of 35 mA (Broser and Luetschg, 2020).

While providing electrical stimuli, a simultaneous trigger signal was sent to the data acquisition system of our MEG system (CTF Omega 275, Coquitlam, BC, Canada). This signal was used to synchronise both the stimulation and the recording. For the recording of the neuromagnetic signal, a sampling rate of 2343.8 samples/s was used. The OPM system delivered two analogue signals for the magnetic field strength in the y and z direction for each sensor. A low-pass filter of 100 Hz was applied to the analogue signal. The OPM system and the analogue digital converter of the recording system both had an intrinsic delay. This delay was measured to be 6.2 ms. The timing of the data was post hoc adjusted so that the stimulus was applied at t = 0 s. During the experiment, 10 electric stimuli were applied, and the evoked magnetic muscular activity was recorded. The plots and analyses of the OPM data was performed in R (R Core Team, Vienna, Austria).

### 2.3 Model of the magnetic field of the MAP

#### 2.3.1 Geometrical considerations

The measurement setup was arranged such that the musculus abductor hallucis was in the xy plane below the OPM sensors T, U and V. Therefore, the position of the MAP on the fibre could be parameterised by:

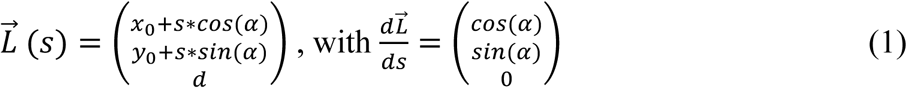

where 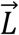(*S*) is the position pointing to the present position of the MAP; *s* is the index to parameterise the longitudinal direction of the muscular fibre; *x*_0_ and *y*_0_ are the translation of the fibre in relation to the OPM sensor in the x and y plane, respectively; *d* is the minimal distance between the OPM sensor and muscle fibre; and *α* is the angle of the muscle fibre in the xy plane defined by the measurement setup.

Without loss of generality the simulation of each sensor was conducted with the assumption that the sensor is located at position (0,0,0). Therefore, for each OPM sensor and each *L*, the point 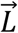(0) corresponded to the point of plumb. Figure 2B shows this geometrical arrangement graphically.

**Figure 2:**
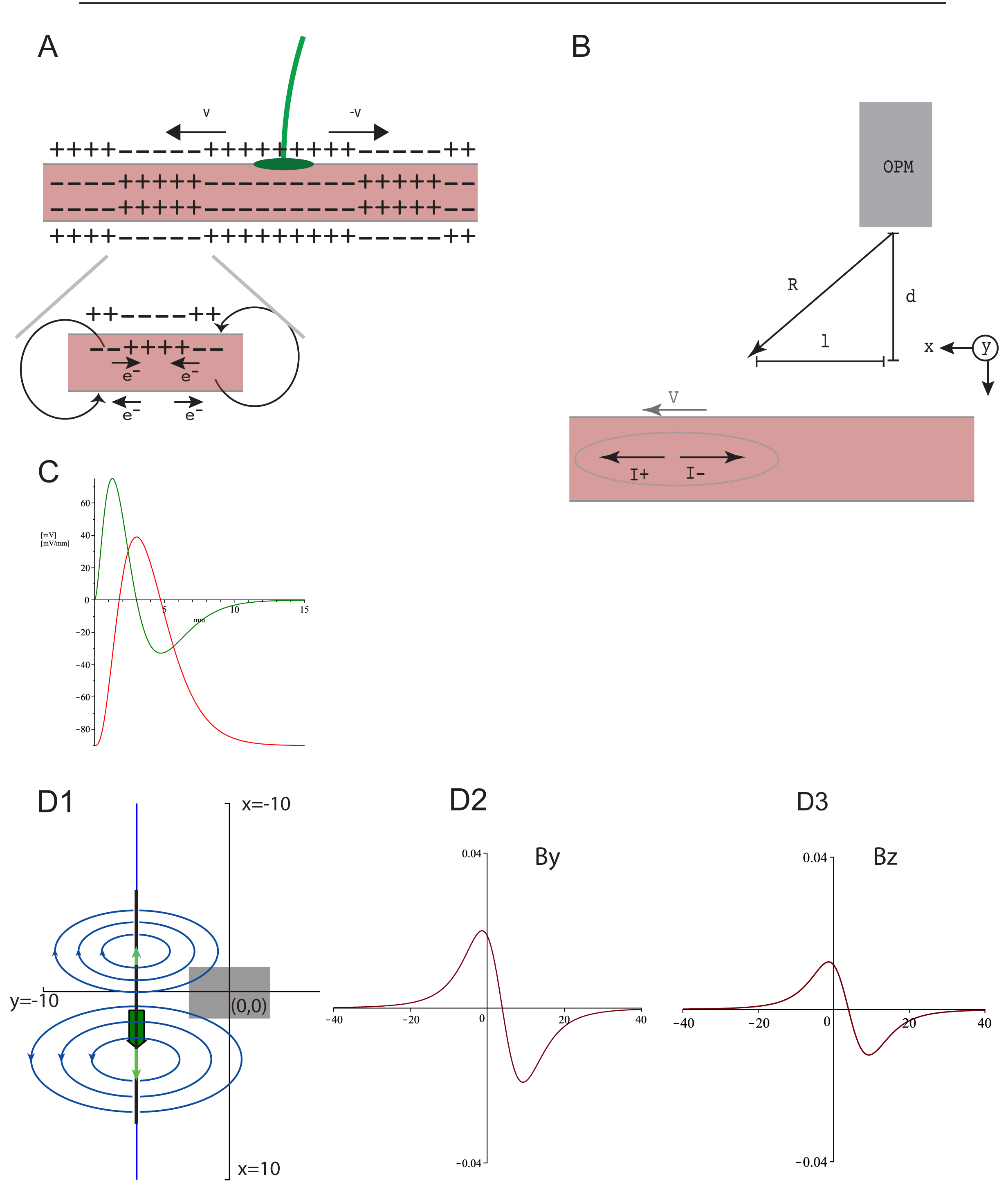
Illustration of the methodological considerations. (A) Schematic drawing of a muscle (dark red) and axon of a motor neuron (green) with the motor endplate. Two (v), (-v) action potentials are shown propagating from the endplate to the distal ends of the muscle fibres. The action potential first generates a current with a technical current direction (movement direction of the positive charges) in the direction of the moving action potential. Then, during the repolarisation phase, a current with a technical current in the opposite direction of the propagating action potential is generated. Both currents generate a circular magnetic field in the direction as defined by the right hand rule. (B) Geometrical situation of OPM device (grey), muscle fibre (dark red) and propagating MAP (grey circle with the currents *I* and movement velocity *V*). In the figure, *d* is the shortest distance between the sensor and muscle, *l* is the longitudinal position of the action potential in relation to the *Lotfußpunkt* (point of shortest distance *d* to the OPM sensor) and *R* is the vector connecting the OPM sensor and position of action potential with the components. The origin of the coordinate system is defined by the position of the OPM sensor.

#### 2.3.2 MAP

According to the work by Farina and Merletti (2001), the voltage at the site of the two propagating MAPs can be described by:

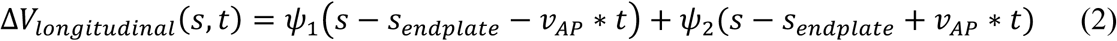

where *ψ*_1_ and *ψ*_2_ being the two MAPs propagating in the proximal and distal directions, respectively. Rosenfalck (1969) described the transmembrane voltage of the MAP using:

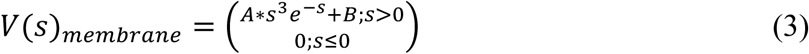

where *A* = 96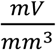 and *B* = −90 mV (Figure 2C). The longitudinal voltage difference can be obtained by differentiation:

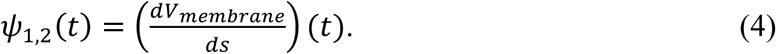

The longitudinal voltage gradient Δ*V*_*longituidnal*_(*s, t*) was divided by the longitudinal resistance *r*_*ix*_ = 173 Ω * *cm* = 17.3 Ω * *mm* (Henneberg and Roberge, 1997) to obtain an estimate of the longitudinal current. The estimation of the current and current density of the radial current in the T-tubuli was more difficult to obtain. Henneberg and Roberge (1997) created a rigorous mathematical approach and values for discovering the radial current. The sum of the currents in the T-tubuli of a given muscular segment is similar in magnitude to the longitudinal current (Henneberg and Roberge, 1997).

#### 2.3.3 Magnetic field of propagating action potentials in general

For isolated nerves, the electric field of a propagating action potential generates a magnetic field (Wikswo et al., 1980; Reincke, 1993) with an absolute magnetic field strength of about 70 pT at a distance of about 1 cm from the nerve surface. Wikswo et al. (1980) showed that when the action potential passes the sensor, a characteristic biphasic magnetic signal can be recorded. The turning point of the signal graph correlates with the action potential below the sensor. In a previous study, we recorded the magnetic field of the muscular activity of the intrinsic hand muscles (Broser et al., 2018). Equation (2) lays the groundwork for the finite wire model to describe the magnetic field generated by the propagating MAP.

### 2.4 Finite wire model

This model assumes that each propagating action potential can be approximated as a finite wire (Figure 2D-1). Without loss of generality, from here on, only one of the two action potentials is considered at a time. The magnetic field of this finite wire can be calculated using the Biot–Savart law (Westgard J., 1997):

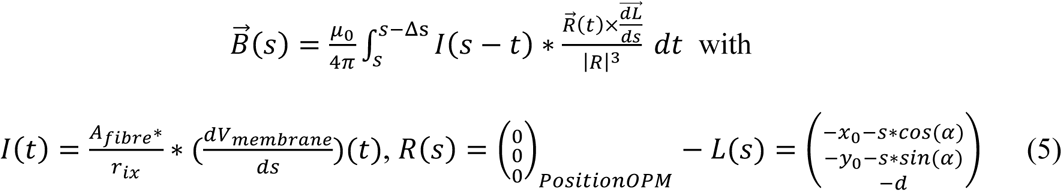

where *s* is the index to parameterise the position of the MAP, Δ*s* = 15 mm and is the longitudinal extent of the MAP (compare with Equation 3 and Figure 2C), *A*_*fibre*_ is the area of electric active tissue in mm^2^, *r*_*ix*_ = 17.3 Ω * *mm* and *I* is the longitudinal current as calculated from Equation (3). The sign of *I* is positive if the movement of positive charges is in the direction of the index parameter *t*.

Remark: *A*_*fibre*_ is dependent on the activated numbers of neuromuscular units are and the total thickness of electric activate muscle tissue.

The experimentally determined parameters were *A*_*fibre*_, *α, y*_*0*_, *x*_*0*_ and *d*. The parameter *d* was estimated by muscle ultrasound to be 8 mm. The computations for the finite wire model were performed using Maple™ (Waterloo, ON, Canada). One simulated example of a resulting magnetic field generated by an action potential propagated according to Figure 2D-1 is shown in Figure 2D-2 in the B field in the y direction and Figure 2D-3 in the B field in the z direction. To quantify the residual relative error of the fit, the *L*^2^ norm on the interval −40 to 40 mm off the residual function 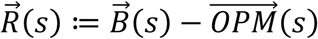 was calculated and normalized by the total signal: 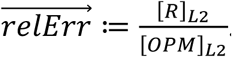.

### 2.5 Magnetic dipole model

Based on our empiric findings, we developed a second model. This model assumes that the dense T-tubuli network in the muscular fibre generates a net radial current. This current then generates a magnetic dipole. Given the finite expansion of the MAP, we hypothesised that this finite magnetic component can be approximated by a magnetic dipole. The magnetic field generated at the site of the OPM sensor would than calculate to:

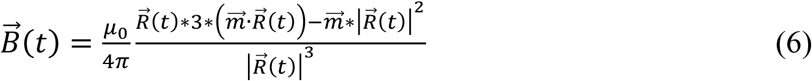

with the dipole vector 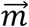 pointing into the direction of the fibre:

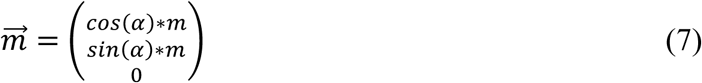

where *m*[*A* · *m*^2^] is the magnitude of the dipole moment. The experimentally determined parameters were *m, α, y*_*0*_, *x*_*0*_ and *d*. Similar to the finite wire model we simulated the propagation of the muscular action potential along the muscular fibre.

## 3 Results

### 3.1 OPM signal

The tibial nerve was stimulated at the level of the knee with 35-mA reliably elicited muscular contractions in the intrinsic foot muscles. As shown in Figure 3, the first signal after stimulation could be detected from the OPM sensors T, U and V about 12 ms after stimulation. In order to estimate the conduction velocity along the tibial nerve, the length of the nerve and the time delay between stimulus onset and the arrival of the signal at the muscle was considered. The lower leg of the subject had a length of 40 cm (knee to medial malleolus), and the distance between the musculus abductor hallucis and the medial malleolus was estimated to be 8 cm. Synaptic transmission typically takes 1 ms. This results in a grossly estimated nerve conduction speed of NCS = (40 cm + 8 cm)/(12 ms - 1 ms) = 0.48 m/11 ms = 43 m/s.

**Figure 3:**
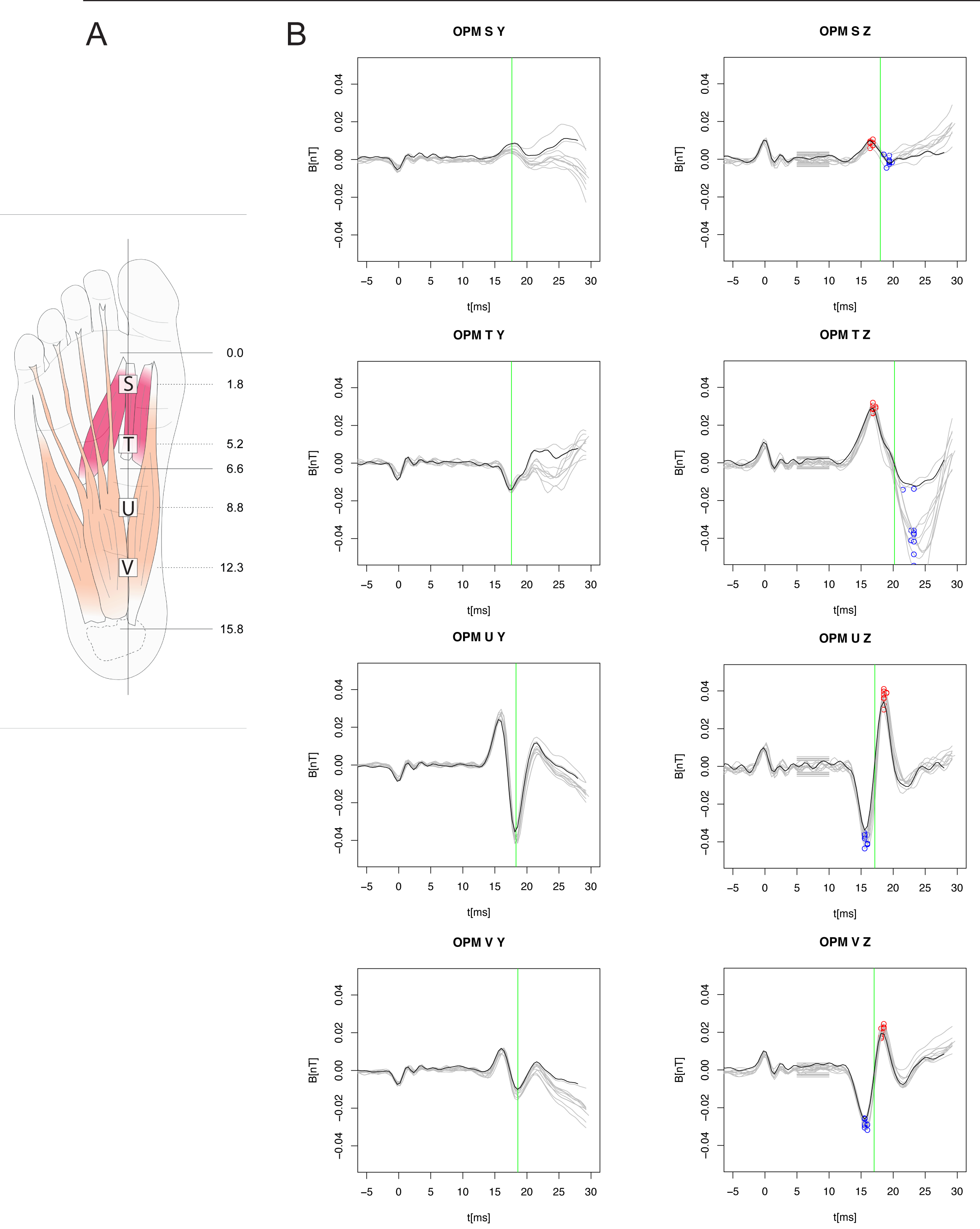
Recorded time-dependent magnetic field strength for each OPM sensor (S, T, U and V). (A) Anatomical location of the four OPM sensors (same as Figure 1). (B) Plots of the recorded time-dependent magnetic field strength in the y (left) and z (right) directions for each sensor S, T, U and V (top to bottom). A total of 10 measurements were performed. Each recorded trace is shown as a fine grey line. The first trace – which was used for further analysis – is shown in black. The small horizontal grey lines shown on the plots for the z direction in the time window 5–10 ms correspond to the minimum/maximum of the signal during this period. These maxima were used to approximate the background signal. The red/blue open circles mark the maximum/minimum of the biphasic peaks. The green line on each graph of the y direction marks the second local maximum/minimum. The green line on the graphs of the z direction marks the turning point of the graph. This point corresponds to the time when the action potential passes the *Lotfußpunkt* (point of shortest distance *d* to the OPM sensor).

A total of 10 stimuli were applied to the nerve. The recorded time-dependent magnetic fields for the sensors in the z and y directions are shown in Figure 3. The signal profiles of the individual traces are quite stable and show only moderate variability. The means and standard deviations of the peak-to-peak amplitude of the signals were 0.0094 ± 0.0021 nT for sensor S, 0.0648 ± 0.0145 nT for sensor T, 0.0759 ± 0.0051 nT for sensor U and 0.0496 ± 0.0029 nT for sensor V. The background signals were 0.0026 ± 0.0007 nT for sensor S, 0.0026 ± 0.0004 nT for sensor T, 0.0038 ± 0.0007 nT for sensor U and 0.0024 ± 0.0006 nT for sensor V. The signal-to-noise ratios were therefore 4:1 for sensor S, 24:1 for sensor T, 20:1 for sensor U and 19:1 for sensor V. Based on the high signal-to-noise ratio, further analysis of the measured data of a single stimulation was conducted. Wikswo et al. (1980) demonstrated that for the propagating axonal action potential, the magnetic flux shows a change in phase while the action potential passes the sensor. The recorded OPM signals showed a similar bi/triphasic profile (Figure 4) with a zero crossing at 18 ms for sensor S, 20 ms for sensor T, 17 ms for sensor U and 17 ms for sensor V. However, given the larger spatial extent of the MAP, the shape of the magnetic flux signal is more complex.

**Figure 4:**
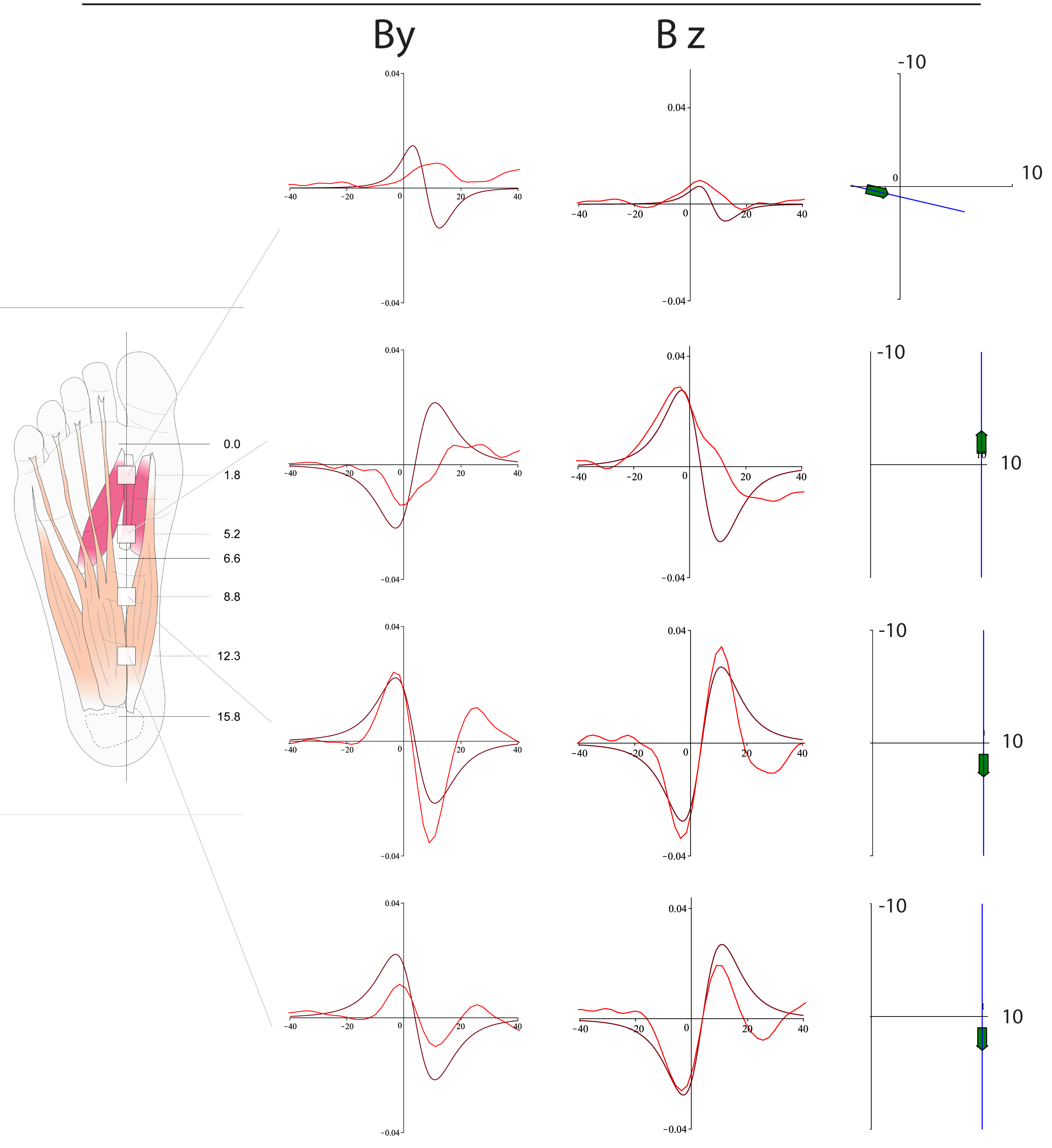
Comparison of experimentally recorded and modelled magnetic field. For each OPM sensor (S, T, U and V), the recorded magnetic field strength in the y and z direction (red trace) is shown. Based on the finite wire model, the hypothetical sources were modelled, and the corresponding magnetic field strengths were plotted (black traces). The plots on the far right show the presumed offset in the y direction, the longitudinal direction in the xy plane of the muscular fibre (black line) and the direction of the MAP. Note the change in the direction of the MAP between sensors T and U. The MAP likely originated between these two sensors.

### 3.2 Finite wire model

In order to account for the complexity of the MAP signal, a manual parameter search was conducted in order to fit the finite wire model onto the data. Sensors T, U and V were anatomically well-located above the musculus abductor hallucis. The muscles beneath sensor S were less well-defined. Therefore, the analysis focused on the sensors T, U and V. By approximating the velocity of the MAP with *v*_*AP*_ = 5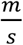 (Farina and Merletti, 2004), we could project the time-depended measured magnetic field strength from the time space into the location space using the formula *z*(*t*) = (*t* − *t*_*offset*_) · *v*_*AP*_. The measurement data were transferred into the location space (Figure 4), and a manual parameter search was conducted with the aim to approximate the strength and signal characteristics.

For each dataset, the model parameters *A*_*fibre*_, *α, y*_*0*_ and *x*_*0*_ were systematically altered, and the signal profiles were visually compared. First the angle *α* was set to α = 0° or α = 180° in order to match the polarity of the first peak in By and Bz direction. Second the parameter *A*_*fibre*_ was screened to obtain a similar amplitude of the signal in the y direction for all three sensors in consideration (T,U,V). Third the *t*_*offset*_ was fine tuned to match the local maxima and minim. Finally, the parameter *y*_*0*_ was varied to match the magnitude of the Bz component. No further parameters had to be adjusted (see Table 1). In order to quantify the goodness of fit the L2 norm of the residual function normalized by the L2 norm of the signal function was calculated (see Table 2).

**Table 1.** Finite wire model predicted parameters

| Sensor | $\alpha$ (°) | $y_0$ (mm) | $A_{fibre}$ (mm<sup>2</sup>) | Offset (ms) |
| --- | --- | --- | --- | --- |
| S | 77 | -4 | 1 | -15.71 |
| T | 180 | 10 | 2 | -17.55 |
| U | 0 | 10 | 2 | -16.30 |
| V | 0 | 10 | 2 | -16.30 |

Note: for all OPM sensors, $x_0 = 0$ and $d = 8$ mm.

**Table 2.** Residual Error - Current Modell

| Sensor | Y – direction | Z – direction |
| --- | --- | --- |
| S | 2.558 | 0.750 |
| T | 2.341 | 0.633 |
| U | 0.375 | 0.287 |
| V | 2.112 | 0.432 |

Interestingly, for sensors T, U and V, only the direction of the propagating action potential (downwards α = 0° or upwards α = 180°) and the time offset (*t*_*offset*_) had to be defined. All other parameters were similar for all three sensors. The electric active fibre area *A*_*fiber*_ was the same for all three sensors.

Based on these parameters, the finite wire model predicted that the MAP propagates beneath sensor T towards the big toe. In addition, it predicted at least one further action potential that propagates in the opposite direction towards the heel and passes sensors U and V. Given that the angles between the two predicted MAPs changed from upwards (*α* _*T*_ = 180°) to downwards (*α*_*U,V*_ = 180°), the MAPs were probably initiated between sensors T and U. This finding directly suggests that the neuromuscular endplate is located between these two sensors.

Comparing the signal traces of the recorded (Figure 4, red traces) and the modelled signals (Figure 4, black traces) reveals a relevant difference that cannot be explained by the finite wire model. In the Y direction, the sensors U and V show a triphasic signal trace. The finite wire model only generates a triphasic signal trace if one considers an obscure geometrical arrangement. Triphasic magnetic signal traces are typical for moving magnetic dipoles.

We therefore empirically tested if a magnetic dipole model (see Supplementary Figure 1) could explain this feature. A parameter search as described above was conducted (Supplementary Table 1) and the signal traces were visualized (see Supplementary Figure 2). As expected, a simulation based on a magnetic dipole returned a triphasic signal trace in the Y direction but a significant angulation of the propagating direction (Supplementary Table 1) was necessary. The residual error for both directions was smaller for the magnetic dipole model when compared to the finite wire model (see Supplementary Table 2).

## 4 Discussion

In this study, we used OPMs to record the magnetic field generated by the propagating action potential of the intrinsic foot muscles. The MAP was triggered by electric stimulation of the supplying nerve in order to elicit synchronised and timely defined MAP of a high number of simultaneously activated neuromuscular units. The study specifically focused on the musculus abductor hallucis brevis. The three sensors located above this muscle picked up the magnetic component of the MAP with a signal that was greater than 10 times the background activity.

As has been previously studied for the nerve AP, we could record a flipping of the B field in the By and Bz direction as the MAP passed the sensor. A manual parameter search was conducted to fit a finite wire model to the data. The only one parameter (*A*_*fiber*_) to be fitted to the data was kept the same for all three sensors in consideration. In addition, based on geometrical anatomical considerations the angle of the propagating direction was selected to be either up or downwards. No further adjustments were necessary. The finite wire model could reasonably reproduce the signal traces and the residual error was measure by the*L*_2_ metric. The error was small for the Bz direction but large for the By direction. Based on this model we were able to predict the location of the neuromuscular endplate.

Although the model explained major features of the recorded data, the triphasic profile of the measured magnetic fields in the By direction could not be explained by the finite wire model. We therefore empirically tested a magnetic dipole model. This model shows the expected signal profiles and the residual errors are smaller in By and Bz direction, but a significant angulation of the dipole would be required which is not supported by the anatomical situation. Therefore, we hypothesize that a combination of the two approaches would model the magnetic field best. However, to do so the recording of the magnetic field in all three spatial directions is required.

### 4.1 Limitations

Even though the study revealed some very interesting findings, it had some relevant limitation. First the OPM sensor used, only recorded the magnetic field in the Z and Y direction. The field strength in X direction could not be measured and a different model might be obtained when the field in this direction would have been recorded.

Second, the study focused on an intrinsic foot muscle. This approach was chosen in order to have a sufficient time delay between electric stimulation and recording. But there are several other muscles in close proximity and so the evaluation of the muscular action potential for the whole extend of the muscle is difficult. In a future study ideally a large muscle with a linear anatomical orientation should be used like the rectus femoris. This muscle would allow to observe the propagation of the muscular action potential over al longer distance. Third, as yet the bandwith of the OPM sensors is limited and a precise temporal and spatial localization of the signal is restricted. Fourth, the study was based on a stimulated approach. That means the supplying nerve was electrically stimulated in order to record the magnetic field. But for a detailed analysis of the neuro-muscular physiology an approach based on the resting activity, the reflex response or the voluntary activity is necessary. This is especially important to establish this method in clinically medicine.

### 4.2 Outlook

We are in the process of planning a new set of experiments using the latest generation of OPM devices to record the magnetic field in all three spatial directions. Given the high signal-to-noise ratio shown by the magnetic component of the MAP, we plan to conduct the next series of experiments based on a voluntary muscular activation paradigm or based on the response of the monosynaptic reflex.

## Abbreviations

OPM: optically pumped magnetometer
MAP: muscle action potential

## Supplementary Tables

**Supplementary Table 1.**
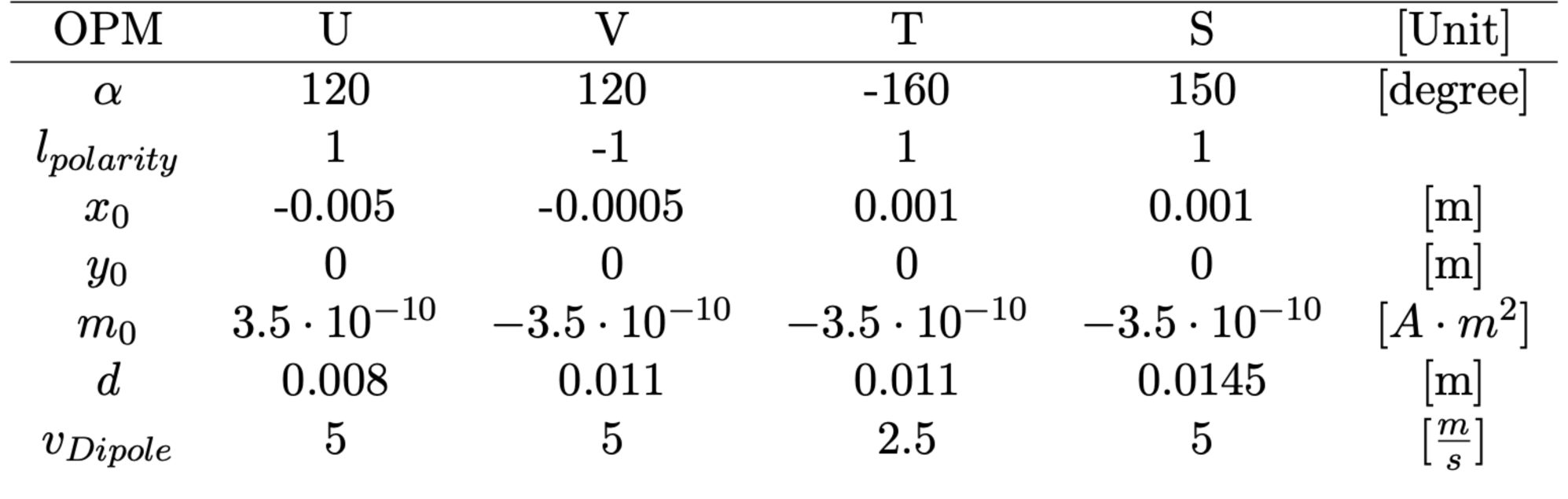
Magnetic dipole model predicted parameters

| OPM | U | V | T | S | [Unit] |
| --- | --- | --- | --- | --- | --- |
| $\alpha$ | 120 | 120 | -160 | 150 | [degree] |
| $l_{polarity}$ | 1 | -1 | 1 | 1 | |
| $x_0$ | -0.005 | -0.0005 | 0.001 | 0.001 | [m] |
| $y_0$ | 0 | 0 | 0 | 0 | [m] |
| $m_0$ | $3.5 \cdot 10^{-10}$ | $-3.5 \cdot 10^{-10}$ | $-3.5 \cdot 10^{-10}$ | $-3.5 \cdot 10^{-10}$ | $[A \cdot m^2]$ |
| $d$ | 0.008 | 0.011 | 0.011 | 0.0145 | [m] |
| $v_{Dipole}$ | 5 | 5 | 2.5 | 5 | $[\frac{m}{s}]$ |

**Supplementary Table 2.** Residual Error - Magnetic Dipole Modell

| Sensor | Y – direction | Z – direction |
| --- | --- | --- |
| S | 0.618 | 1.44 |
| T | 0.428 | 0.464 |
| U | 0.417 | 0.336 |
| V | 1.397 | 0.304 |

---

**Supplementary Figure 1:**
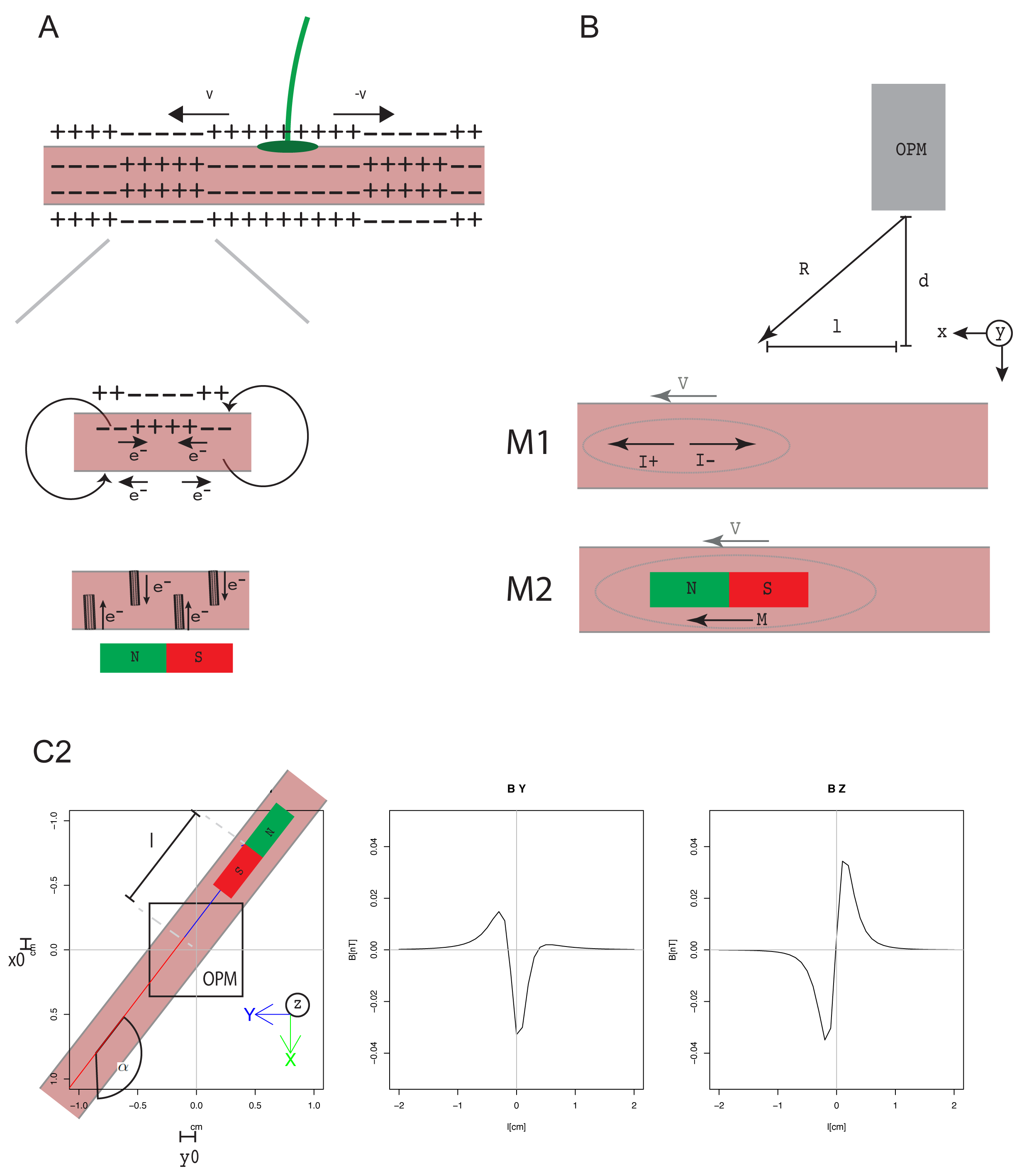
Similar Figure as Figure 2 but with the illustration of the magnetic dipole model. Inset in Panel A illustrates how the radial current along the T – tumuli could generate a magnetic dipole. Graphs in Panel C show the signal profile of a simulated moving magnetic dipole.

**Supplementary Figure 2:**
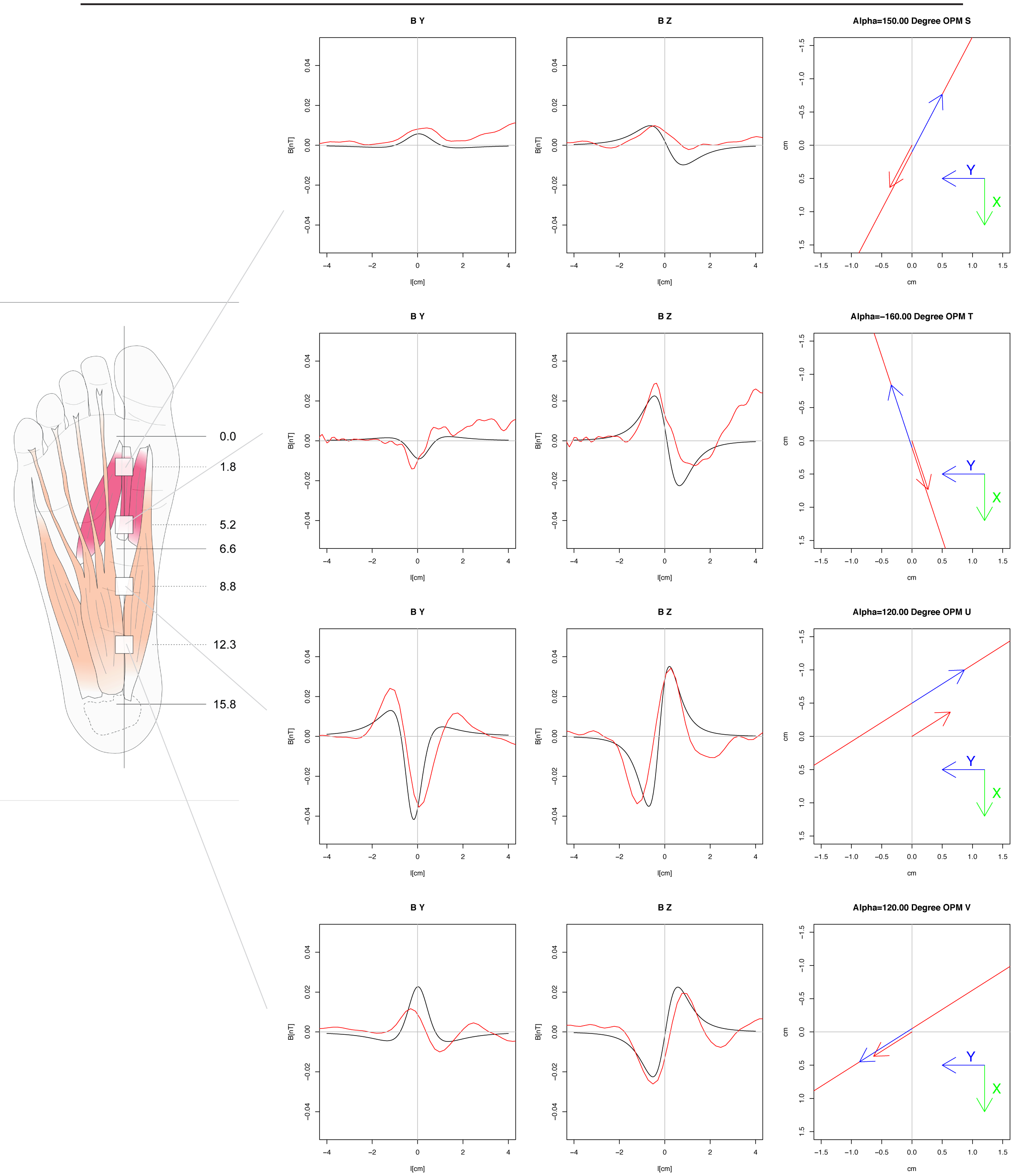
The plot on the left and in the middle show the measured signals for the sensors S, T, U and V top to bottom (red lines). The black line is the result of the simulation based on the magnetic dipole model and model parameters described in the Supplementary Table 1. The axis of abscissae states the distance of the dipole from the “Lotfußpunkt” (point of shortest distance d to the OPM sensor) in cm. The axis of ordinate shows the magnetic field in nano Tesla. The plots on the far right show a two dimensional coordinate system of the xy plane with the z direction point out of the plane and the OPM sensor at the position (x=0,y=0,z=d). The redline on these plots show the linear path of the moving dipole. The blue arrow on the redline points into the direction of movement in relation to time. The red arrow at the centre of origin (x=0, y=0) shows the direction of the magnetic dipole.

